# Chromosomal conservatism vs chromosomal megaevolution: enigma of karyotypic evolution in Lepidoptera

**DOI:** 10.1101/2022.06.05.494852

**Authors:** Elena A. Pazhenkova, Vladimir A. Lukhtanov

**Affiliations:** Department of Biology, Biotechnical Faculty, University of Ljubljana, Večna pot 111, 1000 Ljubljana, Slovenia; Department of Karyosystematics, Zoological Institute of Russian Academy of Sciences, Universitetskaya nab. 1, 199034 St. Petersburg, Russia

**Keywords:** genome, karyotype, chromosomal rearrangement, chromosomal fusion/fission, inversion, translocation, speciation

## Abstract

In the evolution of many organisms, periods of very slow genome reorganization (=chromosomal conservatism) are interrupted by bursts of numerous chromosomal changes (=chromosomal megaevolution). However, the patterns, mechanisms, and consequences of conservative and rapid chromosomal evolution are still poorly understood and widely discussed. Here we show that in blue butterflies (Lepidoptera: Lycaenidae), the periods of chromosome number conservatism are characterized by the real stability of most autosomes and the highly dynamic evolution of the sex chromosome Z, which, due to autosome-sex chromosome fusions and fissions, is carried out according to the cycle Z=>NeoZ_1_=>Z=>NeoZ_2_=>Z=>NeoZ_3_. These fusions and fissions result in a fluctuation of chromosomal number (±1) around the ancestral value, a phenomenon previously observed (but not explained) in numerous groups of Lepidoptera. In the phase of chromosomal megaevolution, the explosive increase in the chromosome number occurs mainly due to simple chromosomal fissions, in some cases complicated by autosomal translocations. Interestingly, these translocations are not random and found to occur only between fragmented chromosomes originated from the same primary linkage group. We also found that the Z chromosomes of two closely related *Lysandra* species are differentiated by a large inversion. We argue that the special role of sex chromosomes in speciation can be reinforced via sex chromosome – autosome fusion. The cycles of fusions and fissions of sex chromosomes with autosomes, such as those found in the blue butterflies, indicate that the species divergence driven by neo-Z chromosome formation is widely distributed in Lepidoptera.

## INTRODUCTION

Chromosomal rearrangements are the most powerful and radical mechanism of genome reorganization in terms of changing the number of chromosomes and the order of genes. Closely related species often differ from each other by structural chromosomal rearrangements, such as chromosome fusions/fissions, translocations, and inversions (White, 1973), but the pattern and evolutionary mechanisms of these changes are not always well understood (Yoshida & Kitano, 2021; Mayrose & Lysak, 2021). One of the mysteries of the structural genome evolution is its extreme temporal irregularity. For many organisms, it is known that periods of very slow genome changes alternate with explosions of chromosomal rearrangements resulting in rapid and complete reshuffling of the linkage groups and gene orders (King, 1993; Wang et al., 2000; Ferguson-Smith & Trifonov, 2007). Examples of such explosive evolution are known for numerous groups of organisms, e.g. mammals (O’Neill et al., 2004), crustaceans (Lécher et al., 1994), and plants (Burchardt et al., 2020). This type of evolution is known as rapid chromosomal change (King, 1993; Mudd et al., 2020), extensive chromosome evolution (Baker & Bickham, 1980), karyotypic megaevolution (Baker & Bickham, 1984; Bell et al., 2001), and runaway chromosomal evolution (Vershinina & Lukhtanov, 2017).

Butterflies and moths (the representatives of the insect order Lepidoptera) are known for showing the most striking examples of both chromosomal conservatism and explosive chromosomal evolution. Most species of Lepidoptera have a haploid number of chromosomes n=31 (Robinson, 1971). This fundamental genome feature, which characterizes the number of linkage groups, is ancestral to the order (Beliajeff, 1930; Suomalainen, 1969) and is preserved in most families for more than 100 million years of their evolution (Lukhtanov 2000, 2014; Ahola et al., 2014). Another example of chromosomal conservatism in Lepidoptera is the preservation of the haploid number n=24 by the majority of the blues butterflies (the family Lycaenidae) (Robinson, 1971). The age of this family was estimated as 78 million years (Espeland et al., 2018).

Despite this general stability, some genera of butterflies and moths show an incredible variety of chromosome numbers (de Lesse, 1960; Robinson, 1971; Brown et al., 2004). For example, in the clade consisting of the genera ((*Polyommatus* +*Neolysandra*)+ *Lysandra*), in less than 5 million years of its evolution (Talavera et al., 2022), a fan of chromosomally differentiated species arose. In these species, the haploid numbers of chromosomes vary from n=10 to n=226 (Lukhtanov et al., 2005; Talavera et al., 2013; Lukhtanov, 2015).

The study of the mechanisms of karyotypic evolution in Lepidoptera was until recently extremely difficult due to the small number of cytogenetic markers suitable for detecting chromosomal rearrangements. Lepidoptera have chromosomes of the holokinetic type, that is, they lack a localized centromere (Murakami & Imai, 1974; Melters et al., 2012; Lukhtanov et al., 2018). This excludes the possibility of using the centromeric index (Levan et al., 1964) as a characteristic of the karyotype. Attempts to use standard differential staining methods such as G- and C-banding of chromosomes have also led to limited success (Bigger, 1975; Goodpasture, 1976).

A significant progress in the study of butterfly karyotypes has been achieved using the FISH and GISH fluorescent staining methods (Šíchová et al., 2015, 2016; Yoshido et al., 2020), but even they do not reveal all the diversity of chromosomal rearrangements that can be expected based on the diversity of chromosome numbers.

The situation has changed quite recently due to the use of chromosome-level genome assemblies for karyotype analysis (Huang et al., 2022; Wang et al., 2022). This led to the first attempts to analyze rapid karyotypic evolution in different groups (Mudd et al., 2020; Yin et al., 2021), including Lepidoptera (Hill et al., 2019; Cicconardi et al., 2021; Mackintosh et al., 2021). However, to the best of our knowledge, no one has studied groups of organisms in which both evolutionary trends (chromosomal conservatism and chromosomal megaevolution) occur simultaneously.

Here we perform analysis of chromosome rearrangements for six species of blue butterflies (Lepidoptera, Lycaenidae). Of these, five species (*Plebejus argus, Cyaniris semiargus, Aricia agestis, Lysandra bellargus*, and *L. coridon*) belong to the *Polyommatus* section (subtribe Polyommatina), and one species (*Glaucopsyche alexis*) belongs to the *Glaucopsyche* section (subtribe Scolitantidini) (Ugelvig et al., 2011; Stradomsky, 2016; Talavera et al., 2022). Four species (*G. alexis, P. argus, C. semiargus*, and *A. agestis*) are characterized by only slightly different haploid chromosome numbers n=23 and n=24 (Robinson, 1971) and represent the evolutionary trend referred to above as chromosomal conservatism. The two closely related species *Lysandra bellargus* and *L. coridon* have highly derived chromosome numbers n=45 and n=90 (Robinson, 1971), respectively. They represent the trend referred to above as chromosomal megaevolution.

## MATERIAL AND METHODS

### Material

Chromosome-level genome assemblies generated by the Darwin Tree of Life Project (https://www.darwintreeoflife.org/) (The Darwin Tree of Life Project Consortium, 2022; Hinojosa Galisteo et al., 2021) and freely available upon deposition in the European Nucleotide Archive (ENA) (https://www.darwintreeoflife.org/wp-content/uploads/2020/03/DToL-Open-Data-Release-Policy-1.pdf) were used for analysis of chromosome structure and detecting chromosome rearrangements (Table S1).

### Detection of macrosynteny and chromosomal rearrangements

We detected macrosynteny in the studied species by pairwise alignment of the assemblies using minimap2 (Li, 2018). To detect the chromosomal rearrangements, the alignments were then visualized as pairwise genomic dotplots with pafr R package (https://github.com/dwinter/pafr). Karyotypes were drawn using RIdeogram R package (Hao et al., 2020).

### Orthologs search and phylogeny reconstruction

The set of chromosome-level genome sequences was screened for core Lepidoptera genes using BUSCO v.5.3.2 (Manni et al., 2021). We found 4899 complete single-copy genes common to all the seven scanned genomes (*Lycaena phlaeas* was used as an outgroup). The protein sequences for each of these genes were obtained for each species and aligned using MAFFT (Katoh & Standley, 2013). A maximum likelihood phylogenetic tree was generated using IQ-TREE 2v with model Q.insect+R5, and node support was estimated using 10,000 bootstrap iterations (Minh et al., 2020).

PAML software (Yang, 2007), specifically, MCMCTree and codeml, were used to estimate divergence dates between studied taxa based on the best obtained ML tree. There is no fossil data available for Lycaenidae, and we used the previously published age of tribe Lycaeninae of ∼60 Mya (Esperand et al., 2018) as a root constrain. The mutation rate for Lepidoptera was previously estimated as 2.9 × 10−9 substitutions per generation (Keightley et al., 2015), equal to 0.725 substitutions per My. We applied an independent clock rate and performed 4 independent runs with 20 million of iterations. Runs were summarized with Logcombiner and parameters were checked in Tracer, which is the part of BEAST package (Bouckaert et al., 2014). The tree was visualised with the Evolview webserver (Subramanian et al., 2019).

### Analysis of repetitive elements

We performed RepeatModeler version 2.0.3 (Flynn et al., 2020) de novo predictions of repetitive elements in six studied chromosome-level assemblies. Resultant libraries were combined using the ReannTE_mergeFasta.pl https://github.com/4ureliek/ReannTE/, which then was used to annotate genomes using RepeatMasker 4 (Smit, 2013-2015).

## RESULTS

### Karyotypes, genome sizes and DNA repeats

According to data from the GenBank (The Darwin Tree of Life Project Consortium, 2022), in the studied species, females are the heterogametic sex; the haploid chromosome numbers (number of autosomes + sex chromosome Z) vary from n=23 to n=90 (Table 1). These numbers are consistent with previously obtained data based on counting the number of chromosomes on cytological preparations (Federley, 1938; Lorkovic, 1941; Bigger, 1960; de Lesse 1960; Robinson, 1971; Talavera et al., 2013).

**Table 1.** Karyotypes and genome sizes of the species studied.

| Species | Haploid chromosome number | Total genome size, Mb | Total size of repeats, Mb | Total size of non-repeat DNA, Mb |
| --- | --- | --- | --- | --- |
| *Glaucopsyche alexis* | n=23 | 639 | 395 | **244** |
| *Plebejus argus* | n=23 | 382 | 181 | **201** |
| *Cyaniris semiargus* | n=24 | 442 | 231 | **211** |
| *Aricia agestis* | n=23 | 435 | 223 | **212** |
| *Lysandra bellargus* | n=45 | 513 | 306 | **207** |
| *Lysandra coridon* | n=90 | 533 | 320 | **213** |

The genomes of the studied species vary greatly in size (from 382 Mb in *Plebejus argus* to 639 Mb in *Glaucopsyche alexis*), and the genome of *Glaucopsyche alexis* is 1.67 times larger than the genome of *Plebejus argus*. We calculated the proportion of repeated elements in the studied genomes (Fig. 1) and chromosomes (Fig. S1), as well as the absolute size of the repeated elements (Table 1) in order to assess the role of the repeated sequences in the detected variations. The analysis showed that the size of non-repeat DNA did not vary significantly in *C. semiargus, A. agestis, L. bellargus* and *L. coridon*, and in general, the differences between the non-repeat parts of the genomes were not so great as the differences between the total genomes. In particular, the total size of non-repeat DNA in *Glaucopsyche alexis* was only 1.2 times larger than the total size of non-repeat DNA in *Plebejus argus*. Thus, the differences in the total genome sizes are determined mainly by repetitive elements.

**Figure 1.**
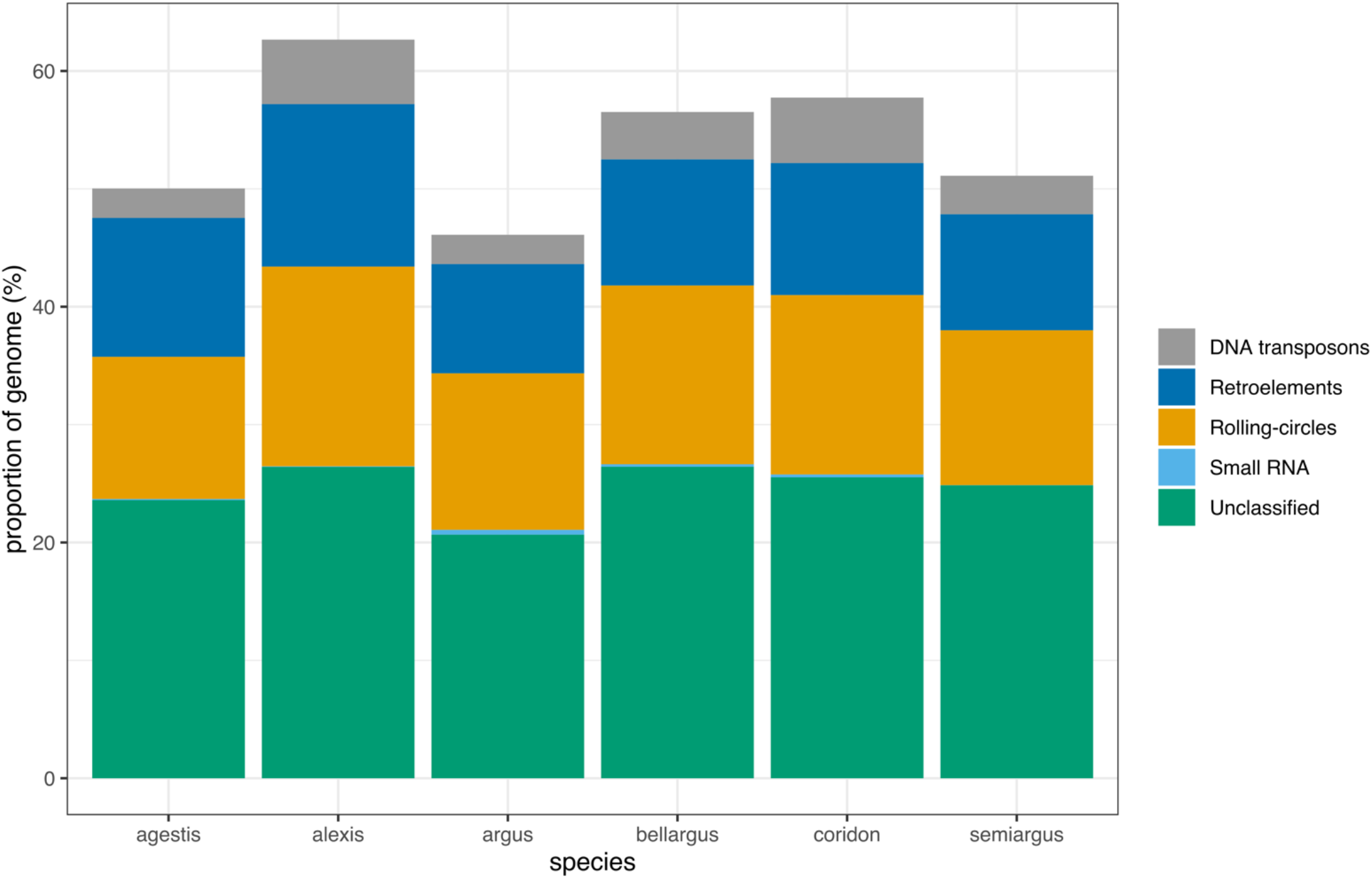
The proportion of the repeatitive elements in the genomes of the studied species.

### Mapping the chromosome numbers on the phylogeny

Phylogenetic analysis of the studied taxa using *Lycaena phlaeas* (n=24; Federley, 1938; Lorkovic, 1941) as an outgroup revealed the topology and dating shown in Fig. 2. The most isolated position on the tree was found in the species *Glaucopsyche alexis*, while the rest of the taxa formed a more compact cluster. This corresponds to the fact that *G. alexis* on one side and the rest of the taxa on the other side belong to different subtribes.

**Figure 2.**
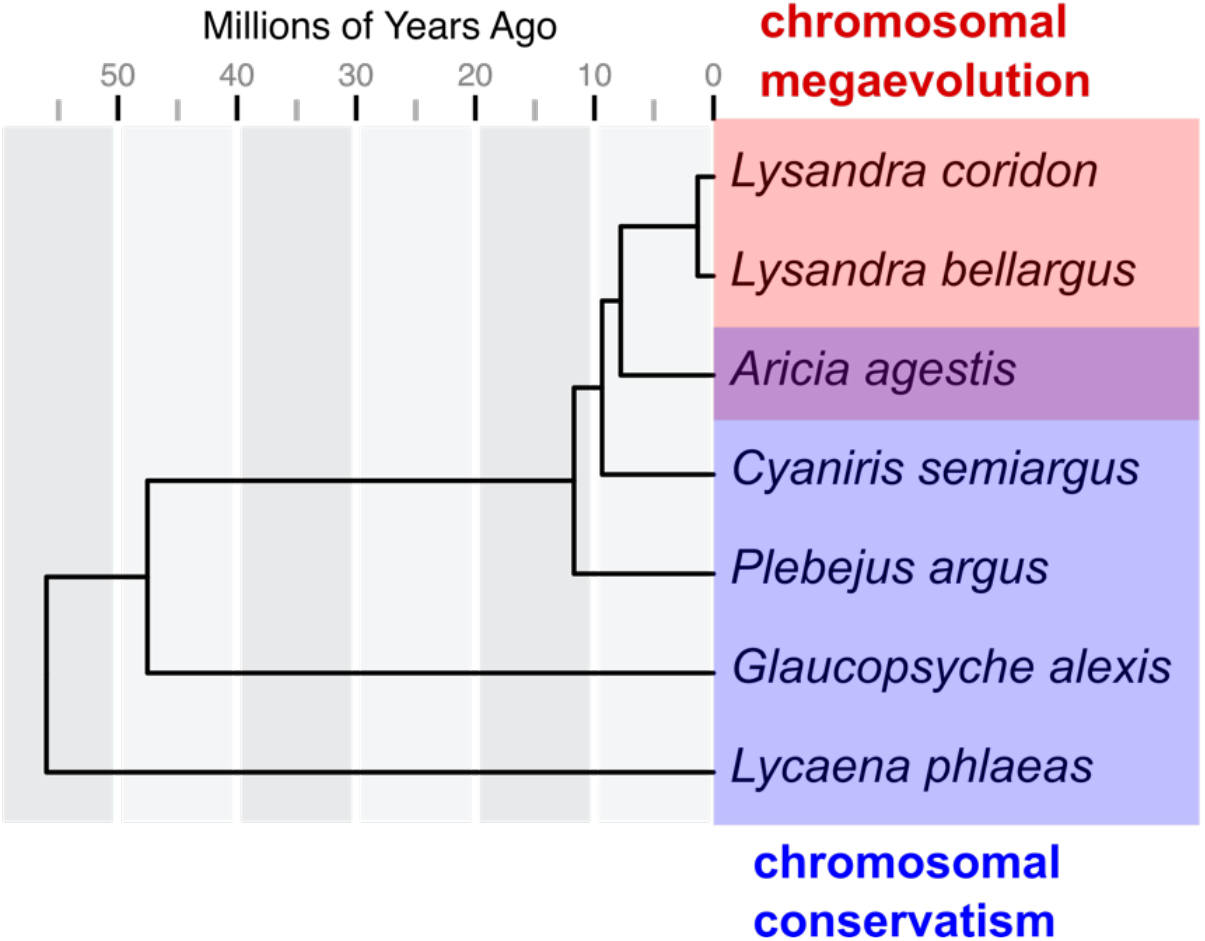
Chromosome numbers of the analyzed taxa mapped on the phylogeny. There are two trends of the chromosome number change: chromosomal conservatism (*Lycaena phlaeas – Glaucopsyche alexis – Plebejus argus – Cyaniris semiargus – Aricia agestis*) (blue) and chromosomal megaevolution (*Aricia agestis – Lysandra bellargus – L. coridon*) (red).

Mapping of the chromosome numbers on the phylogeny (Fig. 2) showed that the taxa *Lycaena phlaeas - Glaucopsyche alexis - Plebejus argus - Cyaniris semiargus - Aricia agestis* represent a conservative phase of karyotypic evolution. During the 55 million years of this phase, chromosome numbers fluctuated between n=23 and n=24. The clade *Aricia agestis - Lysandra bellargus - L. coridon* represents a phase of chromosomal megaevolution. During this phase, chromosome numbers changed from n=23 to n=90 in less than 9 million years.

### Autosomal rearrangements in the phase of chromosomal conservatism

An analysis of species *Glaucopsyche alexis - Plebejus argus - Cyaniris semiargus - Aricia agestis* with “standard” chromosome numbers n=23 and n=24 showed that 20 of their 22-23 autosomes are indeed conservative in the sense that they are not participated in interchromosomal rearrangements (Table 2, Fig. S2). Moreover, a significant proportion of these autosomes were syntenic, that is, they retained the same order of genes. Large inversions that change this order were found only in a part of the chromosomes (Fig. S2). Despite these large scale synteny, homologous chromosomes of these species differ markedly in their genome size (Table S1) which is most likely due to the presence of numerous microinsertions/microdeletions of non-coding sequences.

**Table 2.** Autosomes that did not participate in interchromosomal rearrangements ([although there were inversions in some of them) (green). Autosomes that participated in interchromosomal rearrangements are highlighted by yellow. Each line shows the homologous chromosomes of different species. The serial numbers of chromosomes are presented in accordance with the GenBank (Table S1).

| *Glaucopsyche alexis* (n=23) | *Plebejus argus* (n=23) | *Cyaniris semiargus* (n=24) | *Aricia agestis* (n=23) |
| --- | --- | --- | --- |
| chromosome 1 | chromosome 2 (=1<sub>alexis</sub>) | chromosome 1 (=1<sub>alexis</sub>) | chromosome 1 (=1<sub>alexis</sub>) |
| chromosome 2 | chromosome 3 (=2<sub>alexis</sub>) | chromosome 3 (=2<sub>alexis</sub>) | chromosome 3 (=2<sub>alexis</sub>) |
| chromosome 3 | chromosome 5 (=3<sub>alexis</sub>) | chromosome 4 (=3<sub>alexis</sub>) | chromosome 4 (=3<sub>alexis</sub>) |
| chromosome 4 | chromosome 7 (=4<sub>alexis</sub>) | chromosome 6 (=4<sub>alexis</sub>) | chromosome 6 (=4<sub>alexis</sub>) |
| chromosome 5 | chromosome 4 (=5<sub>alexis</sub>) | chromosome 2 (=5<sub>alexis</sub>) | chromosome 2 (=5<sub>alexis</sub>) |
| chromosome 6 | chromosome 6 (=6<sub>alexis</sub>) | chromosome 5 (=6<sub>alexis</sub>) | chromosome 5 (=6<sub>alexis</sub>) |
| chromosome 7 | chromosome 10 (=7<sub>alexis</sub>) | chromosome 7 (=7<sub>alexis</sub>) | chromosome 7 (=7<sub>alexis</sub>) |
| chromosome 8 | chromosome 8 (=8<sub>alexis</sub>) | chromosome 8 (=8<sub>alexis</sub>) | chromosome 8 (=8<sub>alexis</sub>) |
| chromosome 9 | chromosome 9 (=9<sub>alexis</sub>) | chromosome 9 (=9<sub>alexis</sub>) | chromosome 9 (=9<sub>alexis</sub>) |
| chromosome 10 | chromosome 11 (=10<sub>alexis</sub>) | chromosome 10 (=10<sub>alexis</sub>) | chromosome 10 (=10<sub>alexis</sub>) |
| chromosome 11 | chromosome 12 (=11<sub>alexis</sub>) | chromosome 11 (=11<sub>alexis</sub>) | chromosome 11 (=11<sub>alexis</sub>) |
| chromosome 12 | chromosome 13 (=12<sub>alexis</sub>) | chromosome 12 (=12<sub>alexis</sub>) | chromosome 12 (=12<sub>alexis</sub>) |
| chromosome 13 | chromosome 16 (=13<sub>alexis</sub>) | chromosome 13 (=13<sub>alexis</sub>) | chromosome 13 (=13<sub>alexis</sub>) |
| 14 | 14 (=14<sub>alexis</sub>) | 14 (=14<sub>alexis</sub>) |  |
| chromosome 15 | chromosome 20 (=15<sub>alexis</sub>) | chromosome 19 (=15<sub>alexis</sub>) | chromosome 20 (=15<sub>alexis</sub>) |
| chromosome 16 | chromosome 18 (=16<sub>alexis</sub>) | chromosome 16 (=16<sub>alexis</sub>) | chromosome 15 (=16<sub>alexis</sub>) |
| chromosome 17 | chromosome 15 (=17<sub>alexis</sub>) | chromosome 15 (=17<sub>alexis</sub>) | chromosome 14 (=17<sub>alexis</sub>) |
| chromosome 18 | chromosome 17 (=18<sub>alexis</sub>) | chromosome 17 (=18<sub>alexis</sub>) | chromosome 16 (=18<sub>alexis</sub>) |
| 19 |  |  |  |
| chromosome 20 | chromosome 19 (=20<sub>alexis</sub>) | chromosome 18 (=20<sub>alexis</sub>) | chromosome 17 (=20<sub>alexis</sub>) |
| chromosome 21 | chromosome 22 (=21<sub>alexis</sub>) | chromosome 23 (=21<sub>alexis</sub>) | chromosome 22 (=21<sub>alexis</sub>) |
| chromosome 22 | chromosome 21 (=22<sub>alexis</sub>) | chromosome 22 (=22<sub>alexis</sub>) | chromosome 21 (=22<sub>alexis</sub>) |

### Autosomal rearrangements in the phase of chromosomal megaevolution

The divergence of the clade *Aricia agestis – Lysandra bellargus + L. coridon* was accompanied by a total reorganization of almost all autosomes (Fig. 3). Comparison of *Aricia agestis* with *Lysandra bellargus* shows that these taxa are separated by 18 simple chromosome fissions, one complex fission (chromosome 1 of *Aricia agestis* is divided into three smaller chromosomes 12, 27, and 30 of *L. bellargus*), as well as four major inversions (in chromosomes 6, 21, 28 and 43 of *L. bellargus*) (Fig. S2f). Only three small autosomes of *A. agestis* (19, 21, and 22) did not change.

**Figure 3.**
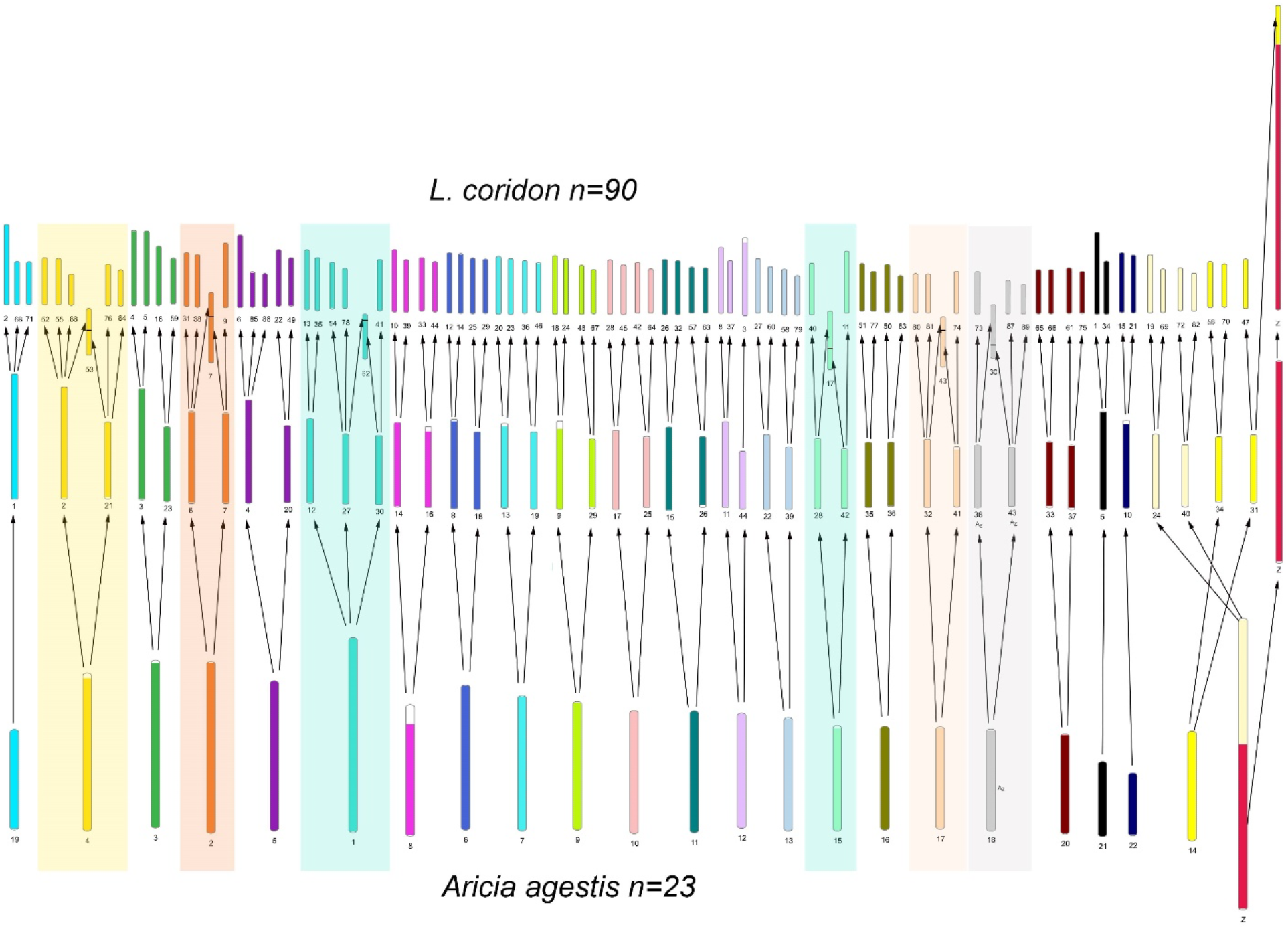
Chromosome megaevolution in the species row *Aricia agestis – Lysandra bellargus – L. coridon*. Karyotypes of *A. agestis* (n=23) (low row) and *L. bellargus* (n=45) (middle row) are differentiated by 22 chromosomal fissions. Karyotypes of *L. bellargus* (n=45) (middle row) and *L. coridon* (n=90) (upper row) are differentiated by 45 fissions, six autosomal translocations and one autosome-sex chromosome translocation. The translocated chromosomes of *L. coridon* (53, 7, 62, 17, 43, and 30) are shown in the figure outside the main chromosome row and are marked with horizontal lines.

Even sharper differences were observed when comparing the closely related species *L. bellargus* and *L. coridon*, separated by only 1.45 million years of evolution. Their comparison showed that only one autosome remained unchanged (chromosome 44_bellargus_ = chromosome 3_coridon_). The karyotypes of these species are separated by 45 fissions, including a set of simple autosomal fissions (one chromosome divides into two) and complex autosomal divisions (one chromosome divides into three), as well as inversions (chromosomes 8, 21, 70, and Z of *L. coridon*). In addition, six translocations were identified (autosomes 7, 17, 30, 43, 53, 62 of *L. coridon*). Interestingly, the autosomal translocations were found to occur only between chromosomes that originated from the same primary linkage group (highlighted by colors in Fig. 3).

### Evolution of the Z chromosome and associated autosomes

The Z chromosome of *Glaucopsyche alexis* is the largest chromosome in the set among the studied species (47.69 Mb). In addition to the core Z sequence, which is also found in the Z chromosomes of all other species studied, it contains a large fragment that is found in autosomes in other studied species (the autosomal A_Z_ fragment) (Fig. 4). Therefore, the Z chromosome of *Glaucopsyche alexis* can be interpreted as a NeoZ chromosome (a result of fusion between the ancestral Z chromosome and an autosome). This interpretation is also supported by the number of chromosomes reduced by one unit (n=23 in *G. alexis*) compared to the modal (and probably ancestral) n=24 (found in *L. phlaeas*).

**Figure 4.**
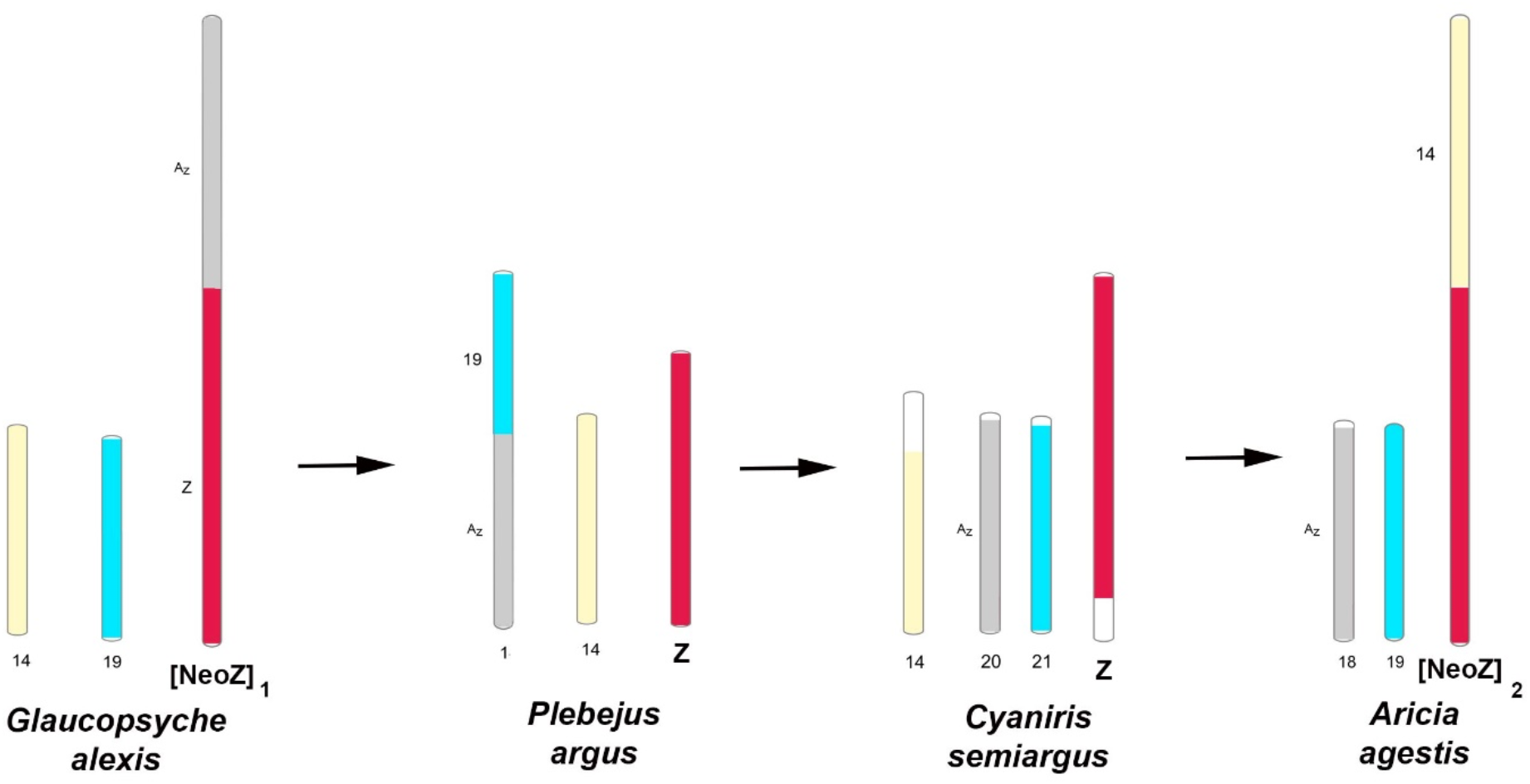
Rearrangements of the Z chromosome and associated autosomes in the phase of chromosomal conservatism. Horizontal arrows indicate row of character transformation (not a direct phylogenetic inheritance of chromosomal characters).

Comparison of Gla*ucopsyche alexis* with *Plebejus argus* shows that the autosomal A_Z_ fragment of the *G. alexis* sex chromosome has translocated to chromosome 19 (Figs 4 and 5). As a result, *Plebejus argus* has a new autosome 1 (= 19_alexis_+ A_Z_), which is the largest element in the chromosome set, and a new chromosome Z.

**Figure 5.**
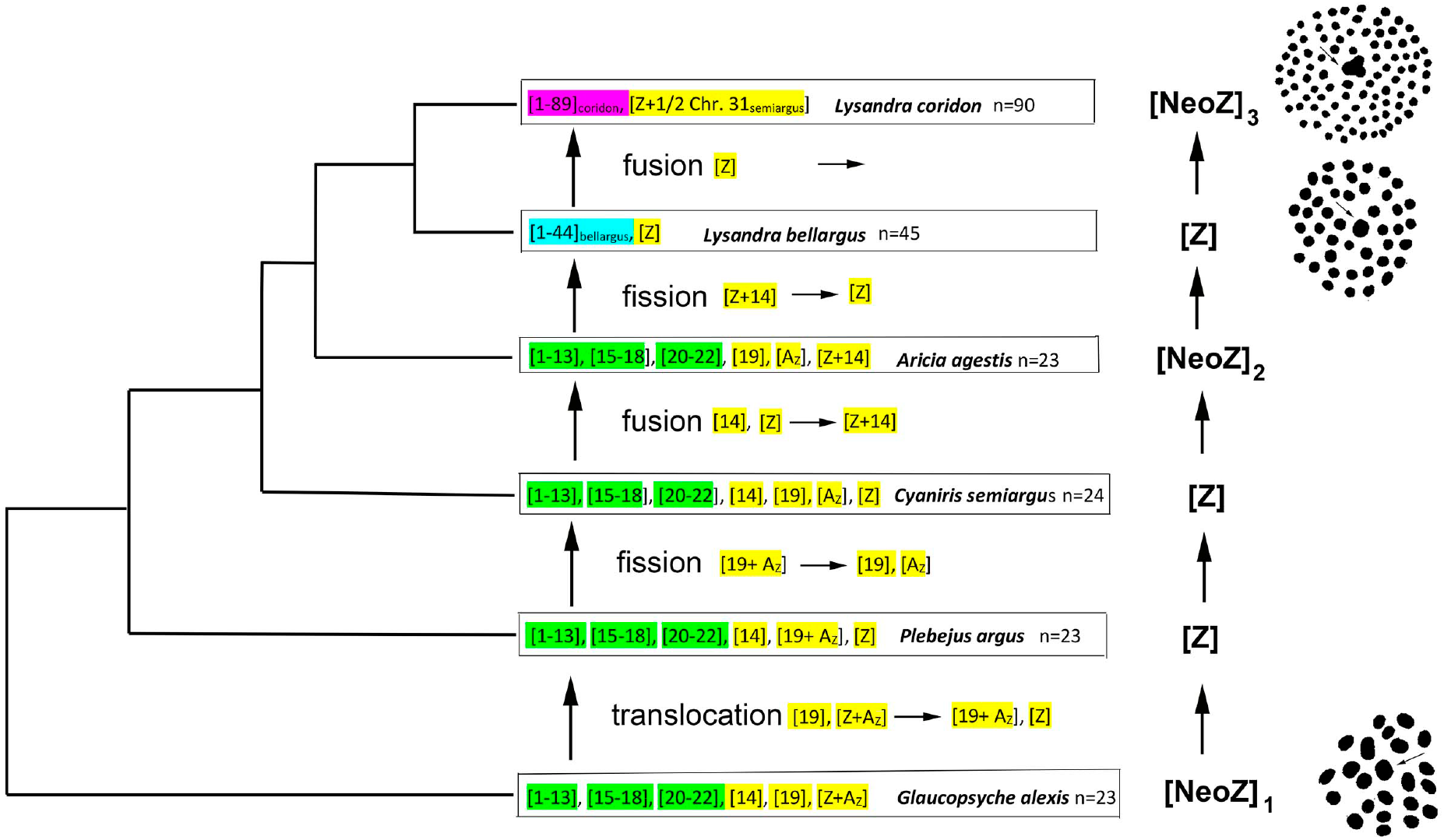
Chromosome rearrangements of the analyzed species mapped on the phylogeny. The conservative chromosomes (not involved in interchromosomal rearrangenrs during the phase of chromosomal conservatism) are shown by green color. The totally rearranged autosomes of *L. bellargus* and *L. coridon* are shown by blue and violet colors, respectively. Z chromosomes and related autosomes are shown by yellow color. Images of metaphase plates at the stage of the first division of male meiosis are given according to Lorkovic (Lorkovic, 1941) (left side of the figure). The largest bivalents (presumably sex Z bivalents) are in the center and marked with arrows. Vertical arrows indicate row of character transformation (not a direct phylogenetic inheritance of chromosomal characters).

When comparing *Plebejus argus* with *Cyaniris semiargus*, it can be seen that the autosome 1 of *Plebejus argus* is divided into two parts (19_alexis_ and A_Z_,), which correspond to the chromosomes 20 and 21 of *Cyaniris semiargus*. As a result, the number of chromosomes in the set increased by one unit.

Comparison of *Cyaniris semiargus* with *Aricia agestis* shows that the fusion of sex chromosome Z with autosome 14 has occurred. As a result, a new sex chromosome (Z + 14) arises; it can be designated as (NeoZ)_2_. The number of chromosomes in the set decreased by one unit.

When comparing *Aricia agestis* with *Lysandra bellargus*, it is seen that the sex chromosome (NeoZ)_2_ divides into three parts resulting in autosomes 24 and 40 and sex chromosome Z of *Lysandra bellargus*. Thus, there is a reversion to the ancestral Z chromosome (Fig. 3).

When comparing *Lysandra bellargus* with *Lysandra coridon*, it can be seen that there was a translocation of the chromosome 31 fragment to the sex chromosome Z (Fig. 3).

### Inversions

After comparisons of pairwise genome-wide alignments, represented as dotplots (Fig. S2), we detected inversions between all the analysed species (Table 3). In the most cases, inversions occurs in autosomes. A large inversion was found in core (non-autosomal) part of Neo-Z sex chromosome in *Lysandra coridon*. Rate of inversion varies in different phylogenetic lineages from 0.125 inversions per My in the pair *G. alexis –P. argus* to 2.7 inverions per My in *L. bellargus – L. coridon*.

**Table 3.**
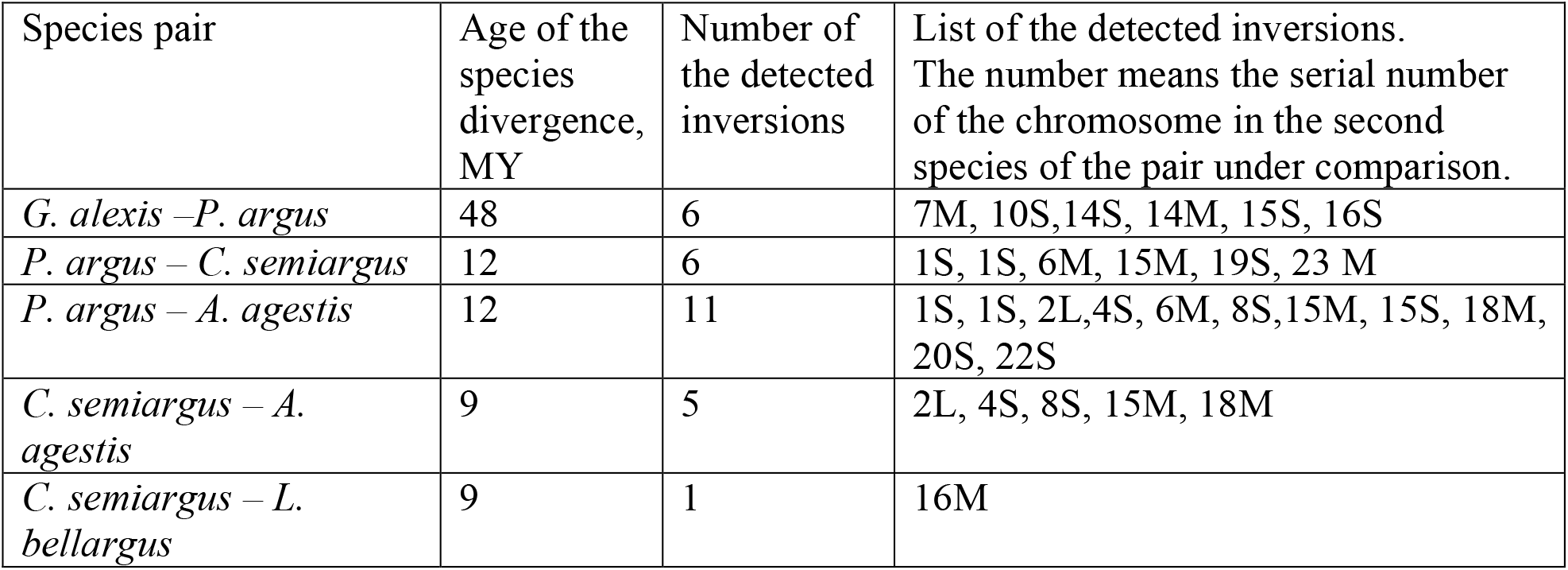
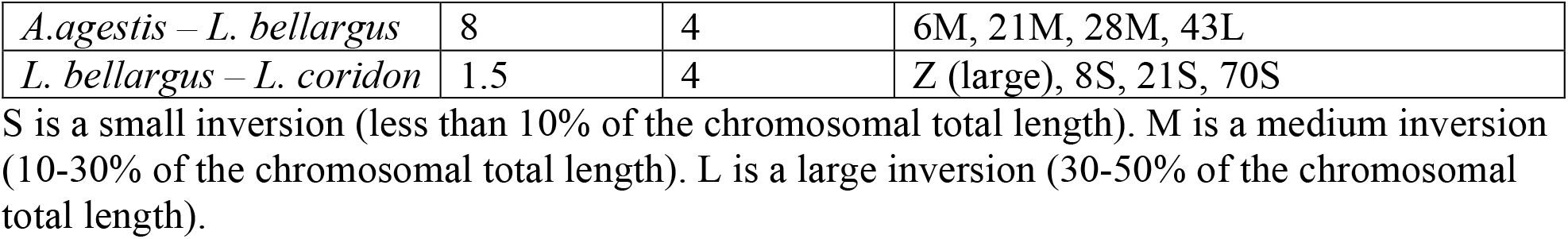
List of the detected inversions.

## DISCUSSION

### Chromosomal conservatism is a real phenomenon

Many researchers, beginning with the pioneering work of Beliajeff (Beliajeff, 1930; Suomalainen, 1969; Robinson, 1971), noted the high conservatism of Lepidoptera karyotypes, based primarily on the stability of chromosome numbers: n=31 for Lepidoptera as a whole (Robinson, 1971), n=25 for the subfamily Pierinae (Lukhtanov, 1992), n=24 for Lycaenidae (Robinson, 1971), n=21 for the genus *Heliconius* (Pringle et al., 2007; Papa et al., 2008; Beldade et al., 2009). However, the stability of chromosome numbers is a weak evidence of the stability of karyotypes, since chromosomal rearrangements, both intrachromosomal (inversions) and interchromosomal (translocations), can lead to a total reorganization of the gene order without changing the number of chromosomes. Indirect evidence in favor of such a mechanism of karyotype evolution was obtained from a comparative analysis of the relative sizes of individual chromosomes in different groups of Lepidoptera with the same number of chromosomes (Lukhtanov and Kuznetsova, 1989). The presence of multiple translocations between species with similar numbers of chromosomes has been directly demonstrated for the family Pieridae (Hill et al., 2019). Despite this, genomic data for other Lepidoptera show a high level of chromosomal synteny when comparing chromosomes of different non-closely related species (Sahara et al., 2007; Pringle et al., 2007; Papa et al., 2008; Beldade et al., 2009; Yasukochi et al., 2009; Ahola et al., 2014; Davey et al., 2016).

Our analysis clearly shows that chromosomal conservatism is a real phenomenon. In the studied group, represented by species *Glaucopsyche alexis - Plebejus argus - Cyaniris semiargus - Aricia agestis*, 20 autosomes out of 22-23 in the set did not participate in interchromosomal rearrangements during more than 45 million years of evolution. Moreover, a significant proportion of these autosomes are syntenic, that is, they retain the same order of genes. Large inversions that change this order were found only in a part (less than half) of the chromosomes.

A high level of macrosynteny does not indicate a complete absence of structural evolution of chromosomes. For several species, the presence of a large number of microarrangements (insertions and deletions) separating homologous chromosomes has been shown (d’Alençon et al., 2010; Ruggieri et al., 2022). Our data also show that against the background of conservation of macrosyntheny (gene order), the compared species differ greatly in the length of DNA repeats. Most likely, these differences arose as a result of microinsertions and microdeletions of the DNA repeats, leading to structural divergence of karyotypes, but not changing the order of genes.

### Chromosomal megaevolution

In contrast to chromosomal conservatism, the phenomenon of rapid karyotypic evolution in the blue butterflies (Lycaenidae) is beyond doubt (Robinson, 1971; Lukhtanov et al., 2005). A comparative phylogenetic analysis of the blue butterfly subgenus *Agrodiaetus* (Lycaenidae: the genus *Polyommatus*), demonstrating a total range of haploid chromosome numbers from n=10 to n=134, revealed a gradual pattern of the rapid chromosome number change (Vershinina & Lukhtanov, 2017). However, the actual cytogenetic mechanisms of this process have not been previously studied.

In our study, a comparison of *Aricia agestis* with *Lysandra bellargus* shows that a rapid twofold increase in the number of chromosomes is achieved through the simplest and most parsimonial way: due to simple chromosome fissions, in which all but three autosomes participate. However, even faster (actually, explosive) chromosome evolution in the lineage leading to *L. coridon* (n=90) shows that the process may be more complex and include, in addition to fissions of most chromosomes, also interchromosomal translocations.

Interestingly, this process, despite its apparent unrestraint, seems to be canalized and not random. This is evidenced by the fact that autosomal translocations were found to occur only between chromosomes that originated from the same primary linkage group. Thus, there is some conservatism even in rapid chromosomal evolution. It manifests itself in the tendency to preserve and re-create the chromosomal sequences that characterized the ancestral linkage groups.

### Evolution of the Z chromosome

The stability of the Z chromosome in Lepidoptera evolution was postulated in early cytogenetic work (White, 1973) and then confirmed in modern studies, which revealed a highly conserved synteny block of ancestral Z-linked genes across Lepidoptera (Dalíková et al., 2017; Fraisse et al., 2017). Our study shows that in the evolution of the Z chromosomes, there is a combination of cytogenetic conservatism of its main core part, apparently inherited from a common distant ancestor of Lepidoptera, with the lability of the autosomal part. This autosomal part changes rapidly in evolution due to a series of successive fusions and/or translocations between the sex chromosomes and different autosomes and subsequent fissions.

The results of fusions between the ancestral sex chromosomes and autosomes are called neo-sex chromosomes (White, 1973; Hejníčková et al., 2021). Neo-Z chromosomes have been repeatedly described as occurring sporadically among different groups of Lepidoptera (Nguyen et al., 2013; Mongue et al., 2017; Mackintosh et al., 2021). More rarely, such sex chromosome – autosome fusions can be fixed in evolution and be the ancestors of entire phylogenetic lineages (Nguyen et al., 2013). Here we describe another, highly dynamic trend of the Z chromosome evolution, which, due to autosome-sex chromosome fusions, is carried out according to the cycle Z=>NeoZ_1_=>Z=>NeoZ_2_=> Z=>NeoZ_3_.

We describe this Z-chromosome cycle for the blue butterflies, but we think that it may be widespread. A sign of this cycle is the fluctuation of chromosome numbers by one unit. It is believed that the chromosome numbers of butterflies are extremely conservative in the vast majority of cases. However, a more careful analysis of the available data shows that the dominant number characteristic of a particular group is usually accompanied by a subdominant one that differs by one unit. So, in nymphalid butterflies (Nymphalidae, subfamily Nymphalinae), the modal n=31 is accompanied by the submodal n=30. In satyrid butterflies (Nymphalidae, subfamily Satyrinae), the modal n=29 is accompanied by the submodal n=28. In the blue butterflies (Lycaenidae), the modal n=24 is accompanied by the submodal n=23 (Robinson, 1971). These subdominant numbers are distributed randomly within these groups, not forming phylogenetic clusters. Our study shows that in the blue butterflies, the fluctuation of chromosomal number (±1) around the ancestral value is explained by autosome – Z chromosome fusions and fissions. We hypothesize that the same mechanism explains the fluctuations in chromosome numbers in other groups of Lepidoptera.

It remains unclear why, against the background of the evolutionary conservatism of autosomes, sex chromosomes show a more dynamic evolution. There are two mutually non-exclusive explanations for this. First, it can be assumed that there are some internal features in the structure of Z chromosomes that predetermine the tendency to merge with autosomes and split. Second, fusions and fissions in the sex chromosomes have a higher chance of being fixed in the population due to the smaller effective population size for the Z chromosome.

### Rapid autosomal fusions/fissions and speciation

Chromosomal heterozygosity leads to the formation of multivalents (instead of normal bivalents) during meiosis. Potentially, this results in segregation problems at the first meiotic division and, as a consequence, complete or partial sterility (Grant, 1981; King, 1993; Borodin et al., 2019). Even a single heterozygous chromosomal rearrangement, such as a reciprocal translocation or chromosomal fusion, is expected to result in 50% reduction of fertility (King, 1993). Therefore, chromosome rearrangements can contribute to the formation of post-zygotic isolation and hence speciation. In fact, the observed number of sterile and/or inviable gametes can be lower than this expectation due to the orientation of multivalents during meiosis (King, 1993), preferential inclusion of inviable nuclei in polar bodies in females (Cortes et al., 2015; Borodin et al., 2019), lower recombination rates in the heterogametic sex (Lenormand and Dutheil, 2005), distorting transmission ratios during meiosis caused by selfish genetic elements (Akera et al., 2019) or inverted meiosis (Lukhtanov et al., 2018). Lepidoptera are known to tolerate heterozygosity for multiple chromosomal rearrangements (Lukhtanov et al., 2011); however, the fertility of these heterozygotes is reduced (Lukhtanov et al., 2018, 2020).). We, therefore, hypothesize that multiple chromosome fissions, such as those seen in *L. bellargus* and *L. corydon*, may contribute to speciation.

### Sex chromosome evolution and speciation

Sex chromosomes evolve faster than autosomes (Charlesworth et al., 2018; Mongue et al., 2022), and, at the same time, in case of interspecific hybridization, are less prone to introgression than autosomes (Martin et al., 2013; Pazhenkova & Lukhtanov, 2021). Both phenomena explain the exceptional role of sex chromosomes in speciation. The fast evolution results in hybrid sterility genes that are preferentially localized on sex chromosomes (Coyne & Orr, 2004; Presgraves, 2008; Cong et al., 2019). The tolerance to introgression prevents incipient species from fusion in case of secondary contact.

Data from different organisms indicate that sex chromosome-autosome fusions can enhance the role of sex chromosomes in speciation and initiate rapid divergence between populations and incipient species (Kitano & Peichel, 2012; Bracewell et al., 2017; Wang et al., 2022). At least three different mechanisms of this process have been described. First, genes that play a role in reproductive isolation between populations might rapidly accumulate on neo-sex chromosomes for many of the same reasons that they are found on ancestral sex chromosomes, thus, promoting speciation (Kitano & Peichel, 2012; Mongue et al., 2022). Second, the fusion of sex chromosomes with autosomes can lead to the formation of new adaptive linkage groups, thereby contributing to ecological speciation (Nguen et al., 2013; Paladino et al., 2019). Third, the sex chromosome – autosome fusions might lead to chromosomal effects affecting reproductive isolation via chromosomal-mediated sterility (King, 1993) and suppression of recombination (Faria & Navarro, 2010). The large scale chromosomal inversions, such as those found in the neo-Z chromosome of *Lysandra coridon*, may be additional barriers (Kirkpatrick & Barton, 2006) contributing to speciation.

Thus, sex chromosomes play a special role in the evolution of reproductive barriers between species, which can be reinforced via sex chromosome – autosome fusion. The cycles of fusions and fissions of sex chromosomes with autosomes, such as those found in the blue butterflies, indicate that the species divergence driven by neo-Z chromosome formation is widely distributed in Lepidoptera.

