## Supplemental Table 1 for "Chromosomal conservatism vs chromosomal megaevolution: enigma of karyotypic evolution in Lepidoptera"

**Table S1. Studied species and chromosomes.**
***Glaucopsyche alexis***
The male *G. alexis* specimen was collected from Alcalá de la Selva, Teruel, Aragon, Spain (latitude 40.3638, longitude -0.7269) (Hinojosa Galisteo et al., 2021).
Genome size (median total length): 639.171 Mb
Median GC%: 36.0499

| **Chromosome** | **GenBank #** | **Size (Mb)** | **GC%** |
|---|---|---|---|
| 1 | FR990043.1 | 39.17 | 35.9 |
| 2 | FR990044.1 | 33.01 | 36.0 |
| 3 | FR990045.1 | 31.81 | 35.9 |
| 4 | FR990046.1 | 31.46 | 36.2 |
| 5 | FR990047.1 | 31.38 | 35.7 |
| 6 | FR990048.1 | 30.88 | 35.8 |
| 7 | FR990049.1 | 29.9 | 36.0 |
| 8 | FR990050.1 | 26.97 | 36.1 |
| 9 | FR990051.1 | 26.52 | 36.0 |
| 10 | FR990052.1 | 25.96 | 35.9 |
| 11 | FR990053.1 | 25.55 | 36.1 |
| 12 | FR990054.1 | 24.58 | 36.1 |
| 13 | FR990055.1 | 23.88 | 36.1 |
| 14 | FR990056.1 | 23.5 | 36.2 |
| 15 | FR990057.1 | 22.56 | 36.4 |
| 16 | FR990058.1 | 22.17 | 36.0 |
| 17 | FR990059.1 | 22.13 | 35.9 |
| 18 | FR990060.1 | 22.05 | 35.9 |
| 19 | FR990061.1 | 22.04 | 36.4 |
| 20 | FR990062.1 | 21.4 | 36.3 |
| 21 | FR990063.1 | 17.19 | 36.6 |
| 22 | FR990064.1 | 16.97 | 36.4 |
| Z | FR990042.1 | 47.69 | 35.3 |

The male *G. alexis* specimen was collected from Alcalá de la Selva, Teruel, Aragon, Spain (latitude 40.3638, longitude -0.7269) (Hinojosa Galisteo et al., 2021).
Hinojosa Galisteo JC, Vila R, Darwin Tree of Life Barcoding collective et al. 2021. The genome sequence of the greenunderside blue, Glaucopsyche alexis (Poda, 1761) [version 1; peer review: awaiting peer review] Wellcome Open Research 2021, 6 :274 https://doi.org/10.12688/wellcomeopenres.17264.1

***Plebejus argus***
Genome size (median total length): 382.014 Mb
Median GC%: 36.6038

| Chromosome | GenBank # | Size (Mb) | GC% |
|---|---|---|---|
| 1 | FR989926.1 | 25.78 | 36.7 |
| 2 | FR989927.1 | 25.11 | 36.9 |
| 3 | FR989928.1 | 20.34 | 36.7 |
| 4 | FR989929.1 | 20.01 | 36.5 |
| 5 | FR989930.1 | 19.93 | 36.7 |
| 6 | FR989931.1 | 19.24 | 36.3 |
| 7 | FR989933.1 | 18.46 | 36.8 |
| 8 | FR989934.1 | 17.02 | 36.7 |
| 9 | FR989935.1 | 16.72 | 36.7 |
| 10 | FR989936.1 | 16.42 | 36.6 |
| 11 | FR989937.1 | 16.34 | 36.1 |
| 12 | FR989938.1 | 16.16 | 36.9 |
| 13 | FR989939.1 | 14.49 | 36.5 |
| 14 | FR989940.1 | 14.38 | 36.6 |
| 15 | FR989941.1 | 14.28 | 36.1 |
| 16 | FR989942.1 | 13.86 | 36.7 |
| 17 | FR989943.1 | 13.56 | 36.5 |
| 18 | FR989944.1 | 13.5 | 36.4 |
| 19 | FR989945.1 | 13.38 | 36.6 |
| 20 | FR989946.1 | 13.02 | 36.9 |
| 21 | FR989947.1 | 9.72 | 37.0 |
| 22 | FR989948.1 | 8.68 | 37.4 |
| Z | FR989932.1 | 18.84 | 36.0 |

***Cyaniris semiargus***
Genome size (median total length): 441.703 MB
Median GC%: 36.4511

| Chromosome | GenBank # | Size (Mb) | GC% |
|---|---|---|---|
| 1 | LR994546.1 | 28.31 | 36.7 |
| 2 | LR994548.1 | 23.83 | 36.3 |
| 3 | LR994549.1 | 23.59 | 36.4 |
| 4 | LR994550.1 | 23.19 | 36.7 |
| 5 | LR994551.1 | 22.67 | 36.3 |
| 6 | LR994552.1 | 21.24 | 36.5 |
| 7 | LR994553.1 | 20.83 | 36.4 |
| 8 | LR994554.1 | 19.76 | 36.6 |
| 9 | LR994555.1 | 19.36 | 36.6 |
| 10 | LR994556.1 | 18.47 | 36.1 |
| 11 | LR994557.1 | 18.09 | 36.7 |

| Chromosome | GenBank # | Size (Mb) | GC% |
|---|---|---|---|
| 12 | LR994558.1 | 17 | 36.4 |
| 13 | LR994559.1 | 16.28 | 36.7 |
| 14 | LR994560.1 | 16.26 | 36.4 |
| 15 | LR994561.1 | 16.13 | 36.0 |
| 16 | LR994562.1 | 15.95 | 36.6 |
| 17 | LR994563.1 | 15.52 | 36.3 |
| 18 | LR994564.1 | 15.2 | 36.5 |
| 19 | LR994565.1 | 14.97 | 36.8 |
| 20 | LR994566.1 | 14.82 | 36.6 |
| 21 | LR994567.1 | 14.58 | 37.0 |
| 22 | LR994568.1 | 11.32 | 36.9 |
| 23 | LR994569.1 | 9.41 | 37.7 |
| Z | LR994547.1 | 24.71 | 35.9 |

***Aricia agestis***
Genome size (median total length): 434.762 Mb
median GC%: 36.8332

| Chromosome | GenBank # | Size (Mb) | GC% |
|---|---|---|---|
| 1 | LR990257.1 | 28.2 | 37.0 |
| 2 | LR990258.1 | 24.84 | 37.0 |
| 3 | LR990259.1 | 24.27 | 36.8 |
| 4 | LR990260.1 | 22.86 | 36.9 |
| 5 | LR990261.1 | 21.7 | 36.6 |
| 6 | LR990262.1 | 21 | 36.9 |
| 7 | LR990263.1 | 19.55 | 36.7 |
| 8 | LR990264.1 | 19.03 | 37.0 |
| 9 | LR990265.1 | 18.58 | 36.9 |
| 10 | LR990266.1 | 17.77 | 36.4 |
| 11 | LR990267.1 | 17.6 | 37.0 |
| 12 | LR990268.1 | 16.64 | 37.1 |
| 13 | LR990269.1 | 16.39 | 36.9 |
| 14 | LR990270.1 | 15.88 | 36.5 |
| 15 | LR990271.1 | 15.27 | 37.0 |
| 16 | LR990272.1 | 15.02 | 36.7 |
| 17 | LR990273.1 | 14.82 | 36.8 |
| 18 | LR990274.1 | 14.66 | 37.1 |
| 19 | LR990275.1 | 14.51 | 37.4 |
| 20 | LR990276.1 | 14.5 | 37.3 |
| 21 | LR990277.1 | 10.87 | 37.7 |
| 22 | LR990278.1 | 9.09 | 37.9 |

| Chromosome | GenBank # | Size (Mb) | GC% |
|---|---|---|---|
| Z | LR990256.1 | 42.15 | 36.2 |

***Lysandra bellargus***
Lysandra bellargus overview
Genome size (median total length): 512.999 Mb
Median GC%: 36.4753

| Chromosome | GenBank # | Size (Mb) | GC% |
|---|---|---|---|
| 1 | HG995320.1 | 18.03 | 36.8 |
| 2 | HG995321.1 | 16.11 | 36.6 |
| 3 | HG995322.1 | 16 | 35.9 |
| 4 | HG995323.1 | 14.92 | 36.0 |
| 5 | HG995324.1 | 14.16 | 36.7 |
| 6 | HG995325.1 | 13.29 | 36.5 |
| 7 | HG995326.1 | 13.03 | 36.4 |
| 8 | HG995327.1 | 12.89 | 36.3 |
| 9 | HG995328.1 | 12.69 | 36.7 |
| 10 | HG995329.1 | 12.63 | 37.0 |
| 11 | HG995330.1 | 12.36 | 36.2 |
| 12 | HG995331.1 | 12.19 | 36.7 |
| 13 | HG995332.1 | 12.19 | 36.4 |
| 14 | HG995333.1 | 12.14 | 36.5 |
| 15 | HG995334.1 | 12.01 | 36.7 |
| 16 | HG995335.1 | 11.66 | 36.6 |
| 17 | HG995336.1 | 11.44 | 35.9 |
| 18 | HG995337.1 | 11.21 | 36.6 |
| 19 | HG995338.1 | 11.11 | 36.5 |
| 20 | HG995339.1 | 11.09 | 36.5 |
| 21 | HG995340.1 | 11.03 | 36.4 |
| 22 | HG995341.1 | 10.81 | 36.6 |
| 23 | HG995342.1 | 10.76 | 37.0 |
| 24 | HG995343.1 | 10.71 | 36.4 |
| 25 | HG995344.1 | 10.24 | 36.2 |
| 26 | HG995345.1 | 10.16 | 36.5 |
| 27 | HG995346.1 | 10.09 | 36.2 |
| 28 | HG995347.1 | 10.09 | 36.4 |
| 29 | HG995348.1 | 10.04 | 36.3 |
| 30 | HG995349.1 | 9.99 | 37.0 |
| 31 | HG995350.1 | 9.86 | 35.9 |
| 32 | HG995351.1 | 9.83 | 36.4 |

| Chromosome | GenBank # | Size (Mb) | GC% |
|---|---|---|---|
| 33 | HG995352.1 | 9.52 | 36.6 |
| 34 | HG995353.1 | 9.46 | 36.2 |
| 35 | HG995354.1 | 9.37 | 36.2 |
| 36 | HG995355.1 | 9.26 | 36.1 |
| 37 | HG995356.1 | 9.25 | 36.7 |
| 38 | HG995357.1 | 8.99 | 36.7 |
| 39 | HG995358.1 | 8.97 | 36.5 |
| 40 | HG995359.1 | 8.94 | 36.6 |
| 41 | HG995360.1 | 8.65 | 36.4 |
| 42 | HG995361.1 | 8.59 | 36.5 |
| 43 | HG995362.1 | 8.41 | 36.3 |
| 44 | HG995363.1 | 8.29 | 36.8 |
| W | HG995364.1 | 4.54 | 36.1 |
| Z | HG995319.1 | 29.06 | 35.9 |

***Lysandra coridon***
Lysandra coridon overview
Genome size (median total length): 532.672 Mb
Median GC%: 36.6328

| Chromosome | GenBank # | Size (Mb) | GC% |
|---|---|---|---|
| 1 | HG992056.1 | 9.18 | 36.7 |
| 2 | HG992057.1 | 9 | 37.5 |
| 3 | HG992058.1 | 8.73 | 36.9 |
| 4 | HG992059.1 | 8.44 | 35.7 |
| 5 | HG992060.1 | 8.34 | 36.5 |
| 6 | HG992061.1 | 8.14 | 36.2 |
| 7 | HG992062.1 | 7.84 | 36.1 |
| 8 | HG992063.1 | 7.39 | 36.6 |
| 9 | HG992064.1 | 7.22 | 36.9 |
| 10 | HG992065.1 | 6.96 | 36.9 |
| 11 | HG992066.1 | 6.92 | 37.1 |
| 12 | HG992067.1 | 6.75 | 36.0 |
| 13 | HG992068.1 | 6.7 | 36.4 |
| 14 | HG992069.1 | 6.69 | 36.8 |
| 15 | HG992070.1 | 6.68 | 37.0 |
| 16 | HG992071.1 | 6.6 | 37.0 |
| 17 | HG992072.1 | 6.57 | 36.4 |
| 18 | HG992073.1 | 6.56 | 37.3 |
| 19 | HG992074.1 | 6.49 | 36.5 |

| Chromosome | GenBank # | Size (Mb) | GC% |
|---|---|---|---|
| 20 | HG992075.1 | 6.47 | 37.1 |
| 21 | HG992076.1 | 6.45 | 37.8 |
| 22 | HG992077.1 | 6.32 | 37.0 |
| 23 | HG992078.1 | 6.26 | 36.3 |
| 24 | HG992079.1 | 6.25 | 36.6 |
| 25 | HG992080.1 | 6.21 | 37.4 |
| 26 | HG992081.1 | 6.18 | 37.3 |
| 27 | HG992082.1 | 6.09 | 36.3 |
| 28 | HG992083.1 | 6.07 | 36.2 |
| 29 | HG992084.1 | 6.06 | 36.2 |
| 30 | HG992085.1 | 6.03 | 36.6 |
| 31 | HG992086.1 | 6.03 | 36.1 |
| 32 | HG992087.1 | 6.02 | 36.5 |
| 33 | HG992088.1 | 5.98 | 36.7 |
| 34 | HG992089.1 | 5.95 | 37.1 |
| 35 | HG992090.1 | 5.93 | 37.6 |
| 36 | HG992091.1 | 5.89 | 36.9 |
| 37 | HG992092.1 | 5.87 | 36.3 |
| 38 | HG992093.1 | 5.87 | 37.4 |
| 39 | HG992094.1 | 5.86 | 36.3 |
| 40 | HG992095.1 | 5.77 | 36.9 |
| 41 | HG992096.1 | 5.72 | 37.7 |
| 42 | HG992097.1 | 5.66 | 36.5 |
| 43 | HG992098.1 | 5.65 | 36.1 |
| 44 | HG992099.1 | 5.61 | 36.8 |
| 45 | HG992100.1 | 5.57 | 36.1 |
| 46 | HG992101.1 | 5.54 | 36.5 |
| 47 | HG992102.1 | 5.54 | 35.9 |
| 48 | HG992103.1 | 5.47 | 37.0 |
| 49 | HG992104.1 | 5.41 | 36.2 |
| 50 | HG992105.1 | 5.37 | 36.3 |
| 51 | HG992106.1 | 5.3 | 36.8 |
| 52 | HG992107.1 | 5.27 | 37.8 |
| 53 | HG992108.1 | 5.24 | 36.1 |
| 54 | HG992109.1 | 5.21 | 36.4 |
| 55 | HG992110.1 | 5.2 | 37.1 |
| 56 | HG992111.1 | 5.15 | 36.1 |
| 57 | HG992112.1 | 5.13 | 36.4 |
| 58 | HG992113.1 | 5.13 | 37.0 |
| 59 | HG992114.1 | 5.13 | 38.3 |

| Chromosome | GenBank # | Size (Mb) | GC% |
|---|---|---|---|
| 60 | HG992115.1 | 5.12 | 37.3 |
| 61 | HG992116.1 | 5.07 | 36.6 |
| 62 | HG992117.1 | 5.05 | 36.4 |
| 63 | HG992118.1 | 5.03 | 37.3 |
| 64 | HG992119.1 | 4.93 | 36.0 |
| 65 | HG992120.1 | 4.92 | 36.5 |
| 66 | HG992121.1 | 4.91 | 37.2 |
| 67 | HG992122.1 | 4.91 | 36.3 |
| 68 | HG992123.1 | 4.85 | 36.7 |
| 69 | HG992124.1 | 4.85 | 37.1 |
| 70 | HG992125.1 | 4.83 | 36.7 |
| 71 | HG992126.1 | 4.79 | 36.4 |
| 72 | HG992127.1 | 4.77 | 36.8 |
| 73 | HG992128.1 | 4.76 | 37.0 |
| 74 | HG992129.1 | 4.69 | 36.7 |
| 75 | HG992130.1 | 4.68 | 37.4 |
| 76 | HG992131.1 | 4.67 | 36.3 |
| 77 | HG992132.1 | 4.46 | 36.0 |
| 78 | HG992133.1 | 4.45 | 36.6 |
| 79 | HG992134.1 | 4.42 | 36.3 |
| 80 | HG992135.1 | 4.37 | 36.3 |
| 81 | HG992136.1 | 4.34 | 36.9 |
| 82 | HG992137.1 | 4.22 | 37.1 |
| 83 | HG992138.1 | 4.12 | 36.9 |
| 84 | HG992139.1 | 4.03 | 37.3 |
| 85 | HG992140.1 | 3.94 | 36.3 |
| 86 | HG992141.1 | 3.74 | 36.0 |
| 87 | HG992142.1 | 3.63 | 36.7 |
| 88 | HG992143.1 | 3.49 | 36.4 |
| 89 | HG992144.1 | 3.17 | 36.7 |
| Z | HG992055.1 | 34.01 | 36.1 |
