## Supplemental Figure 1 for "Chromosomal conservatism vs chromosomal megaevolution: enigma of karyotypic evolution in Lepidoptera"

### Supplementary figures

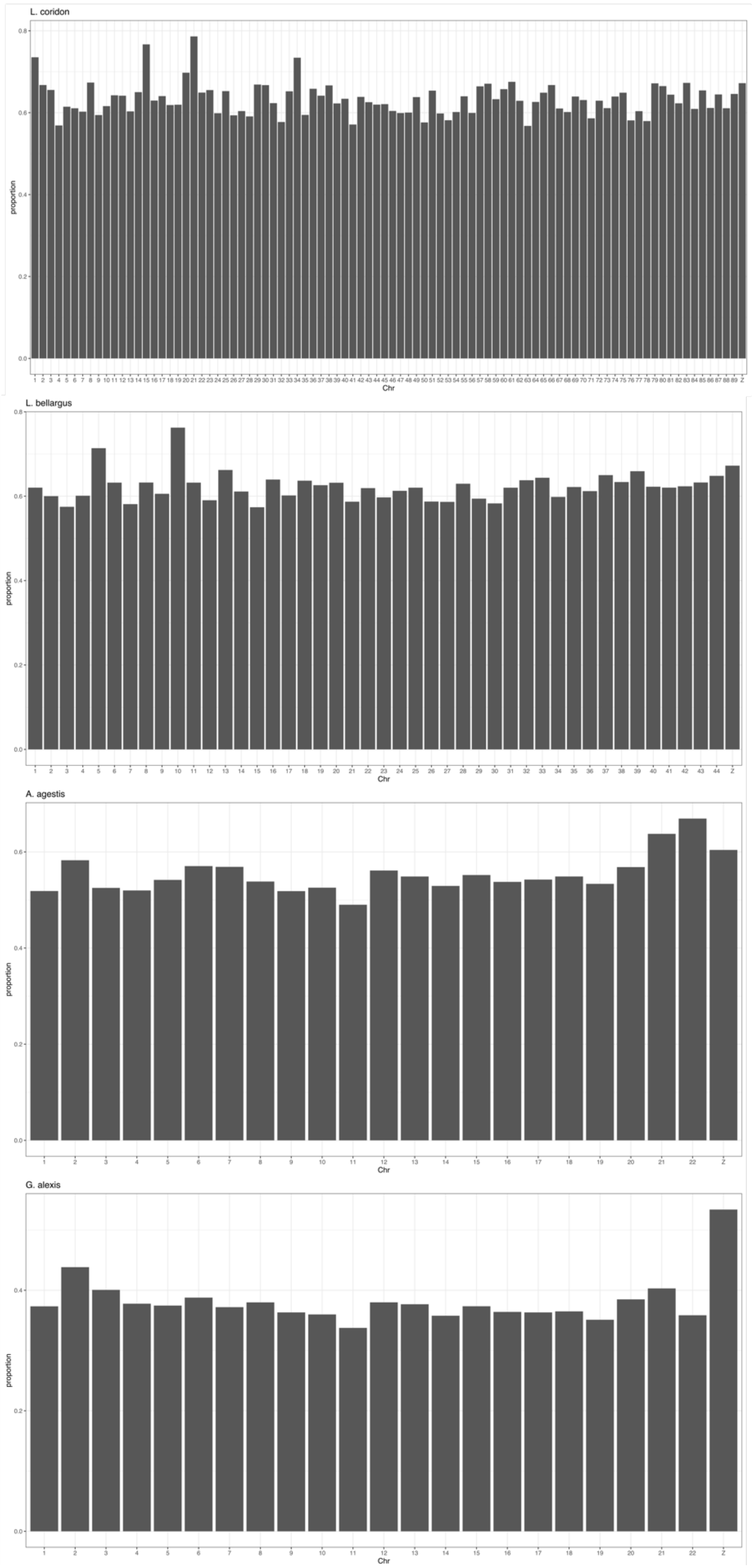


**Figure S1. Proportion of repetitive elements by chromosome in *Lysandra coridon*, *L. bellargus*, *Aricia agestis* and *Glaucopsyche alexis* genomes.**
