## Supplemental Figure 2 for "Chromosomal conservatism vs chromosomal megaevolution: enigma of karyotypic evolution in Lepidoptera"

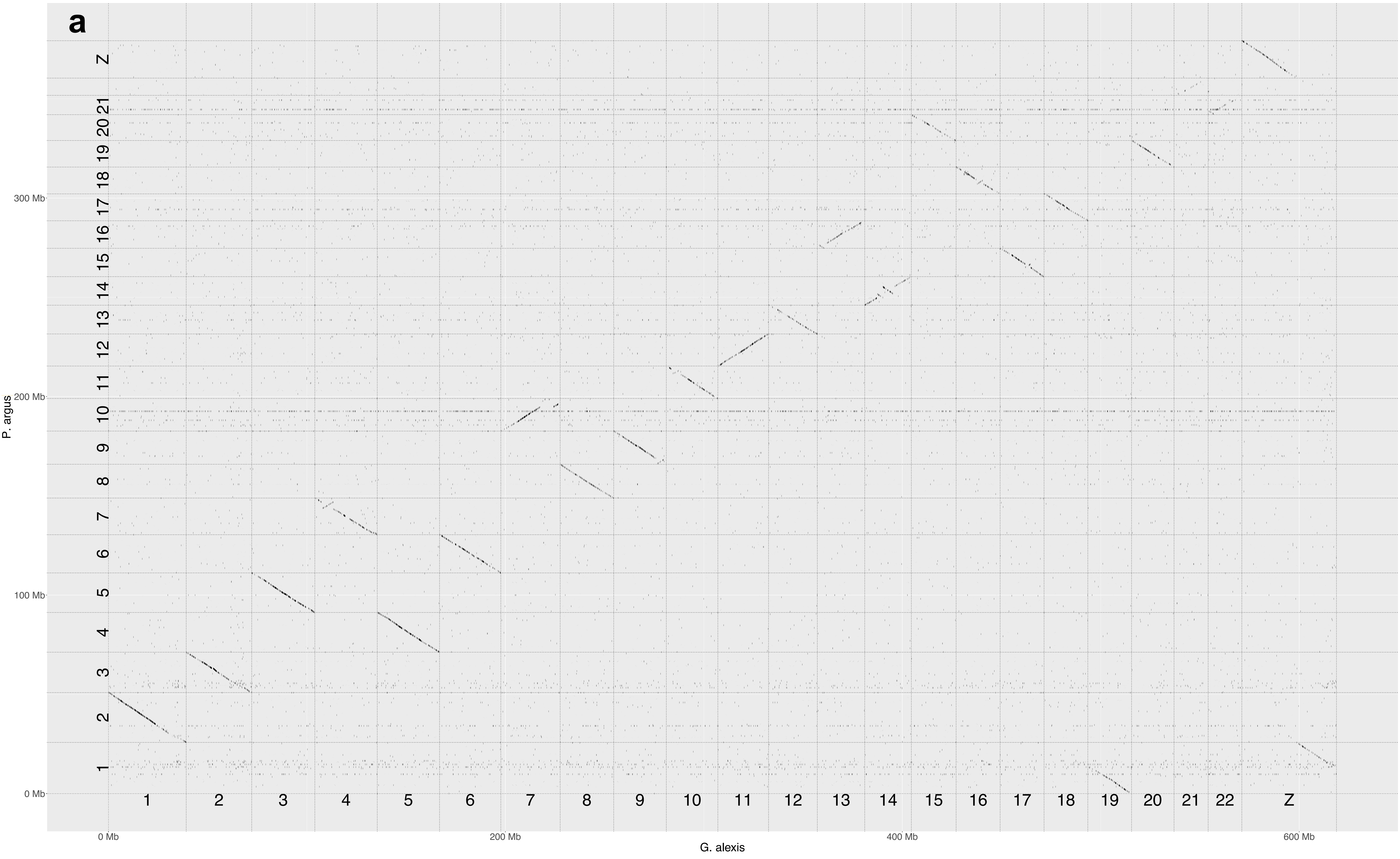

**Figure S2. Genome-wide dot plot for pairwise alignment between studied species.**

**a.** *Glaucopsyche alexis - Plebejus argus*, **b.** *Cyaniris semiargus - P. argus*, **c.** *Aricia agestis - C. semiargus*, **d.** *A. agestis - P. argus*, ***e.*** *Lysandra bellargus - C. semiargus*, **f.** *L. bellargus - A. agestis*, **g.** *Lysandra coridon - L. bellargus*

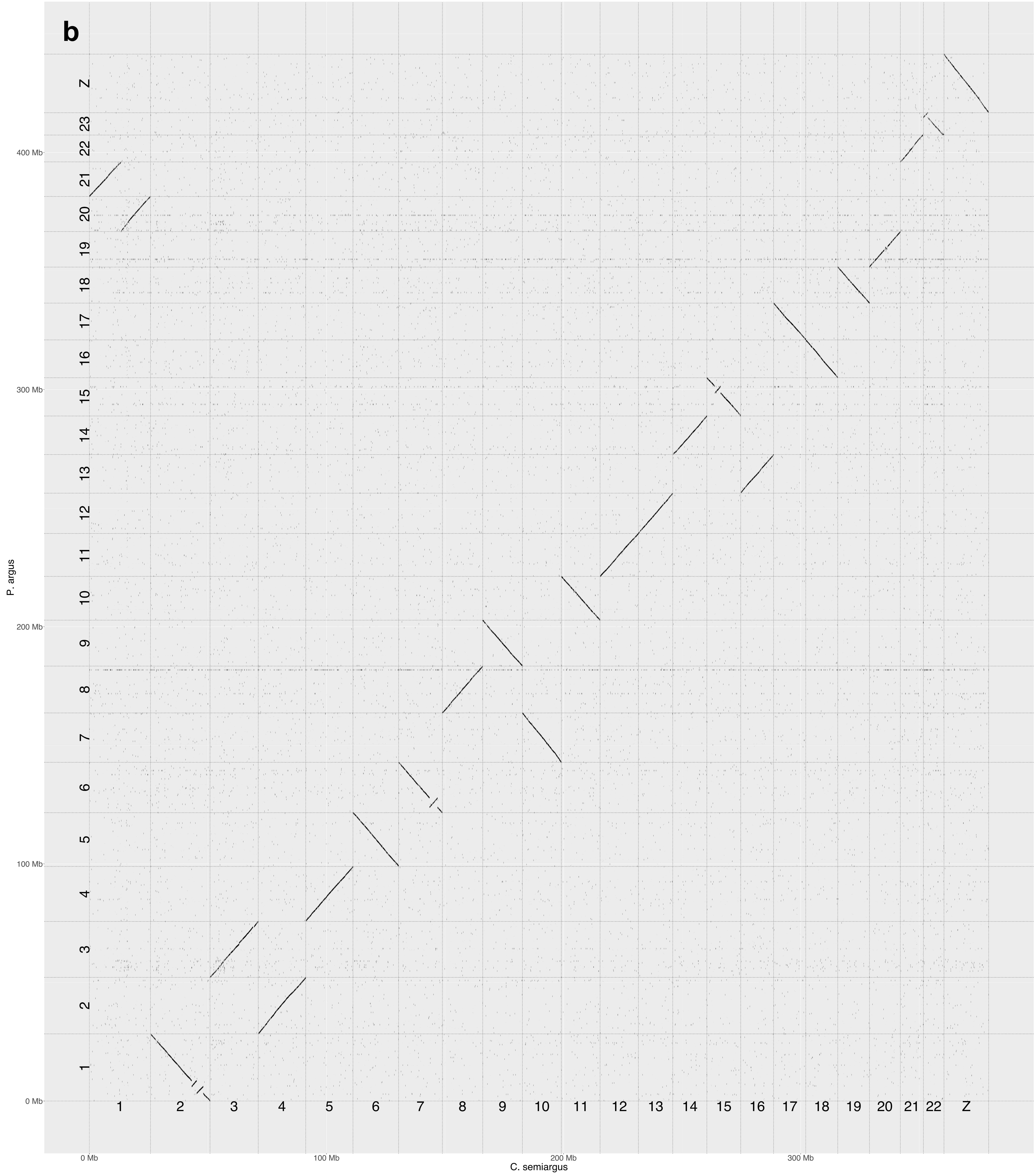

b
P. argus
C. semiargus
0 Mb
100 Mb
200 Mb
300 Mb
400 Mb
Z

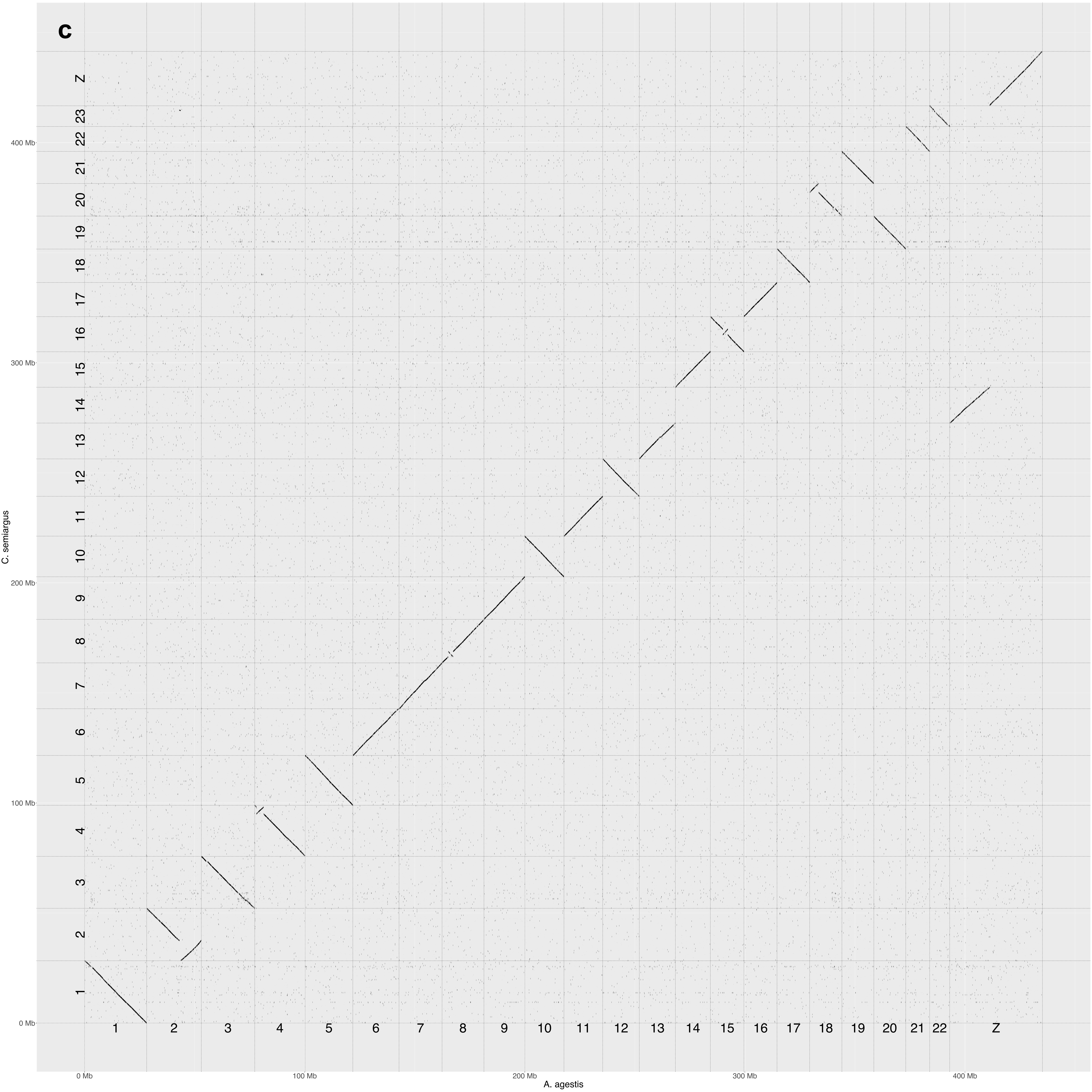
c
Z
23
22
21
20
19
18
17
16
15
14
13
12
11
10
9
8
7
6
5
4
3
2
1
400 Mb
300 Mb
200 Mb
100 Mb
0 Mb
C. semiargus
Z
0 Mb
100 Mb
200 Mb
300 Mb
400 Mb
A. agestis

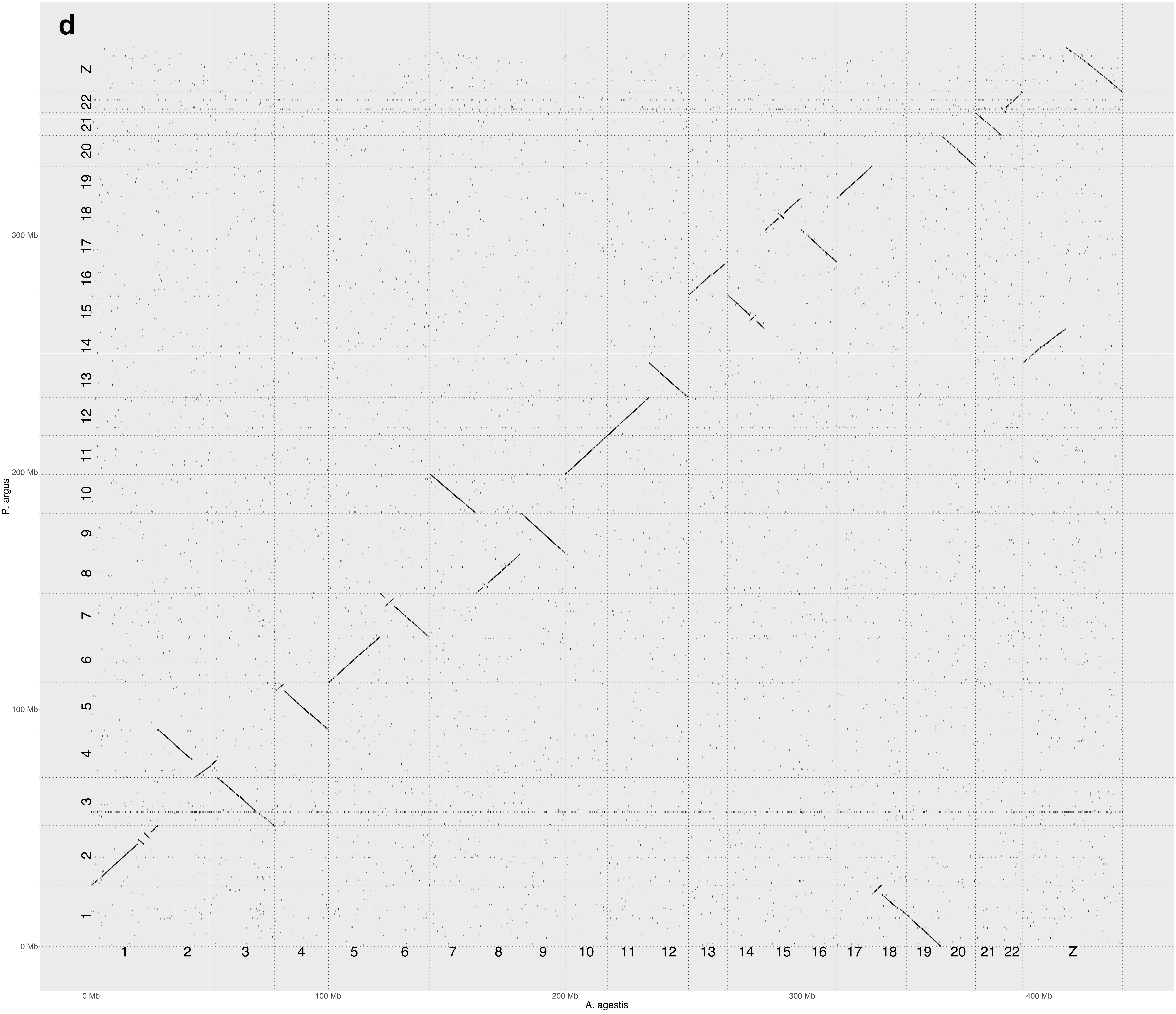
d
P. argus
A. agestis
0 Mb
100 Mb
200 Mb
300 Mb
400 Mb
1 2 3 4 5 6 7 8 9 10 11 12 13 14 15 16 17 18 19 20 21 22 Z

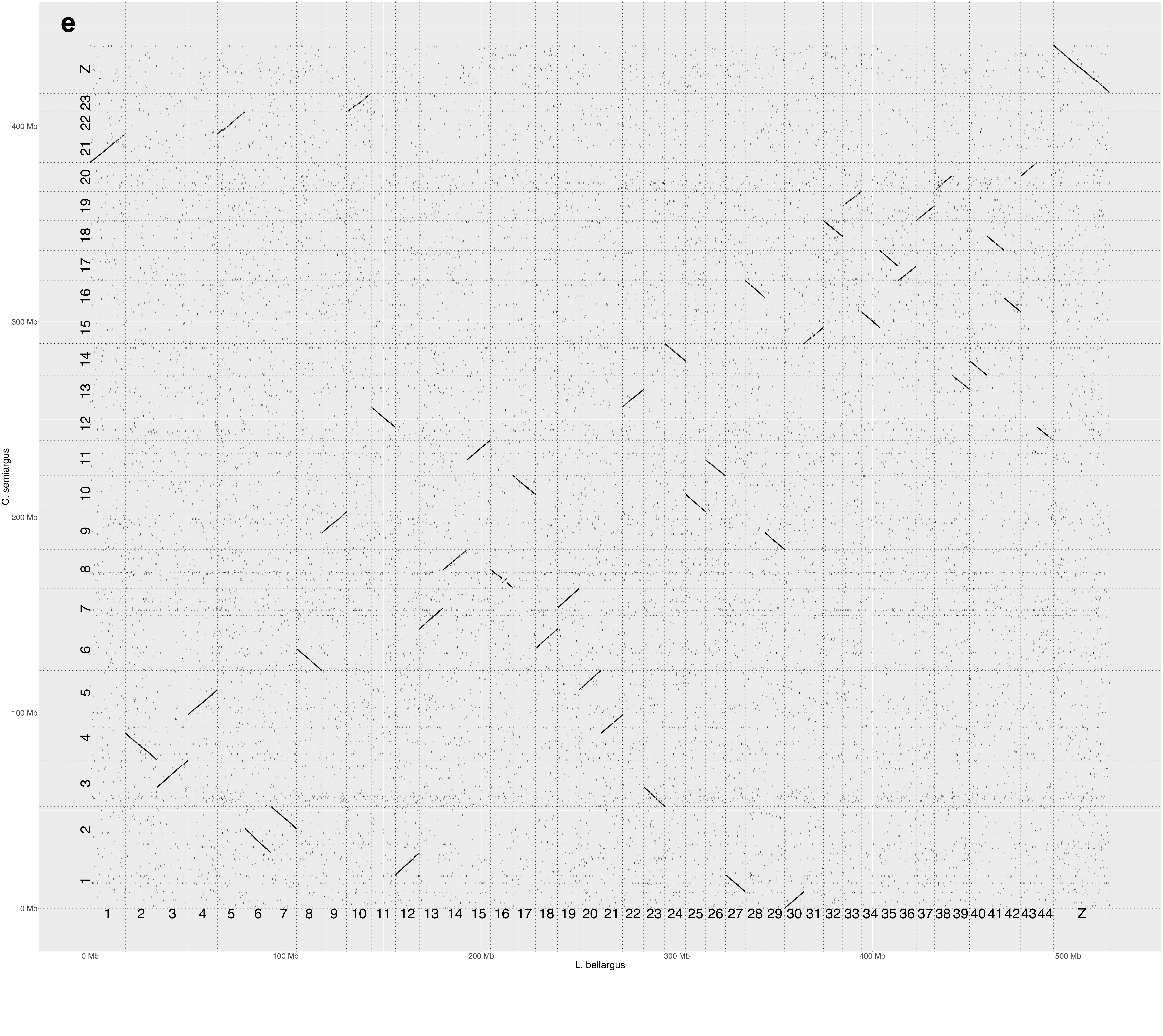

e
C. semiargus
Z
23
22
21
20
19
18
17
16
15
14
13
12
11
10
9
8
7
6
5
4
3
2
1
400 Mb
300 Mb
200 Mb
100 Mb
0 Mb
1 2 3 4 5 6 7 8 9 10 11 12 13 14 15 16 17 18 19 20 21 22 23 24 25 26 27 28 29 30 31 32 33 34 35 36 37 38 39 40 41 42 43 44 Z
0 Mb
100 Mb
200 Mb
300 Mb
400 Mb
500 Mb
L. bellargus

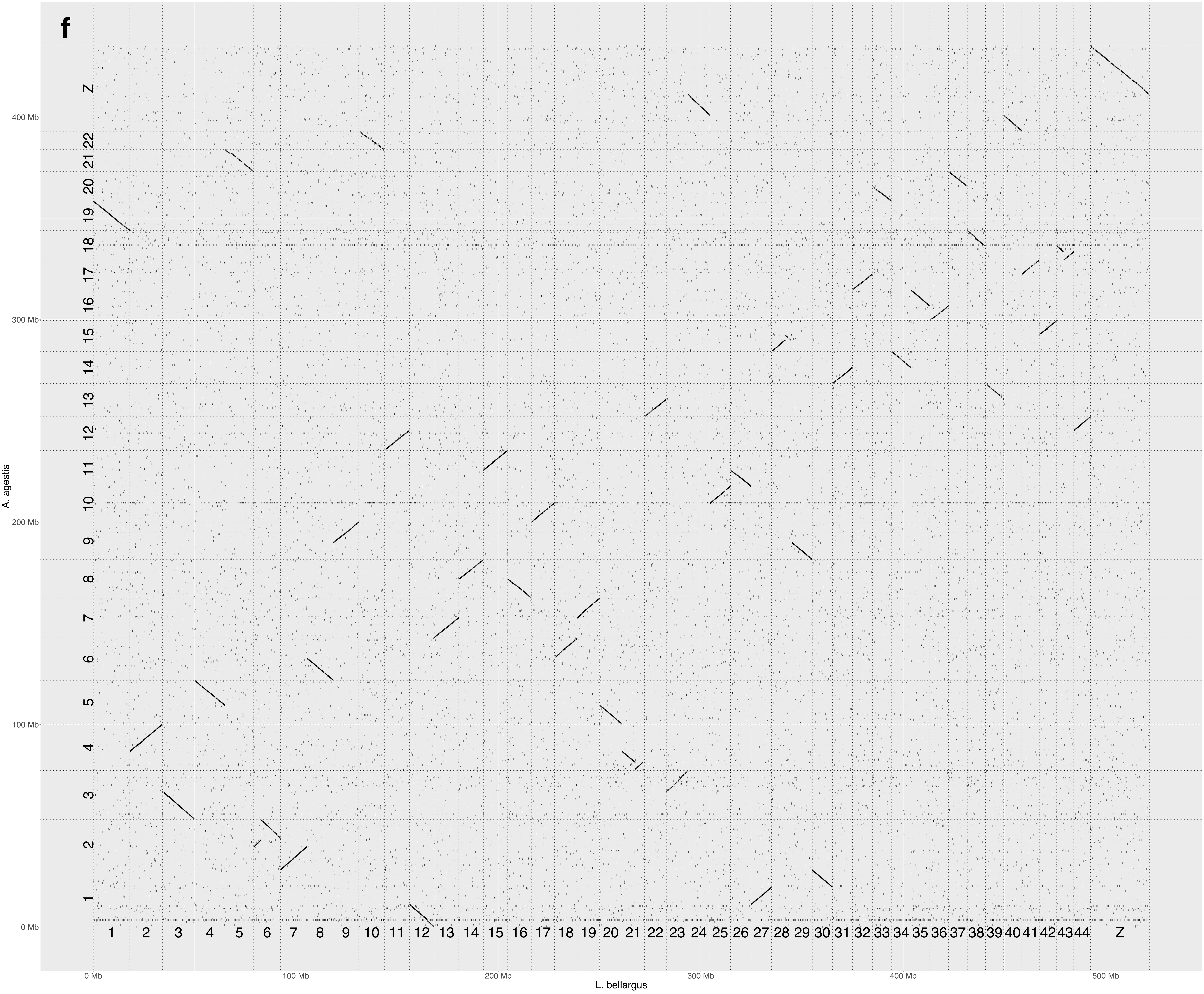
f
Z
22
21
20
19
18
17
16
15
14
13
12
11
10
9
8
7
6
5
4
3
2
1
400 Mb
300 Mb
200 Mb
100 Mb
0 Mb
A. agestis
1 2 3 4 5 6 7 8 9 10 11 12 13 14 15 16 17 18 19 20 21 22 23 24 25 26 27 28 29 30 31 32 33 34 35 36 37 38 39 40 41 42 43 44 Z
0 Mb
100 Mb
200 Mb
300 Mb
400 Mb
500 Mb
L. bellargus

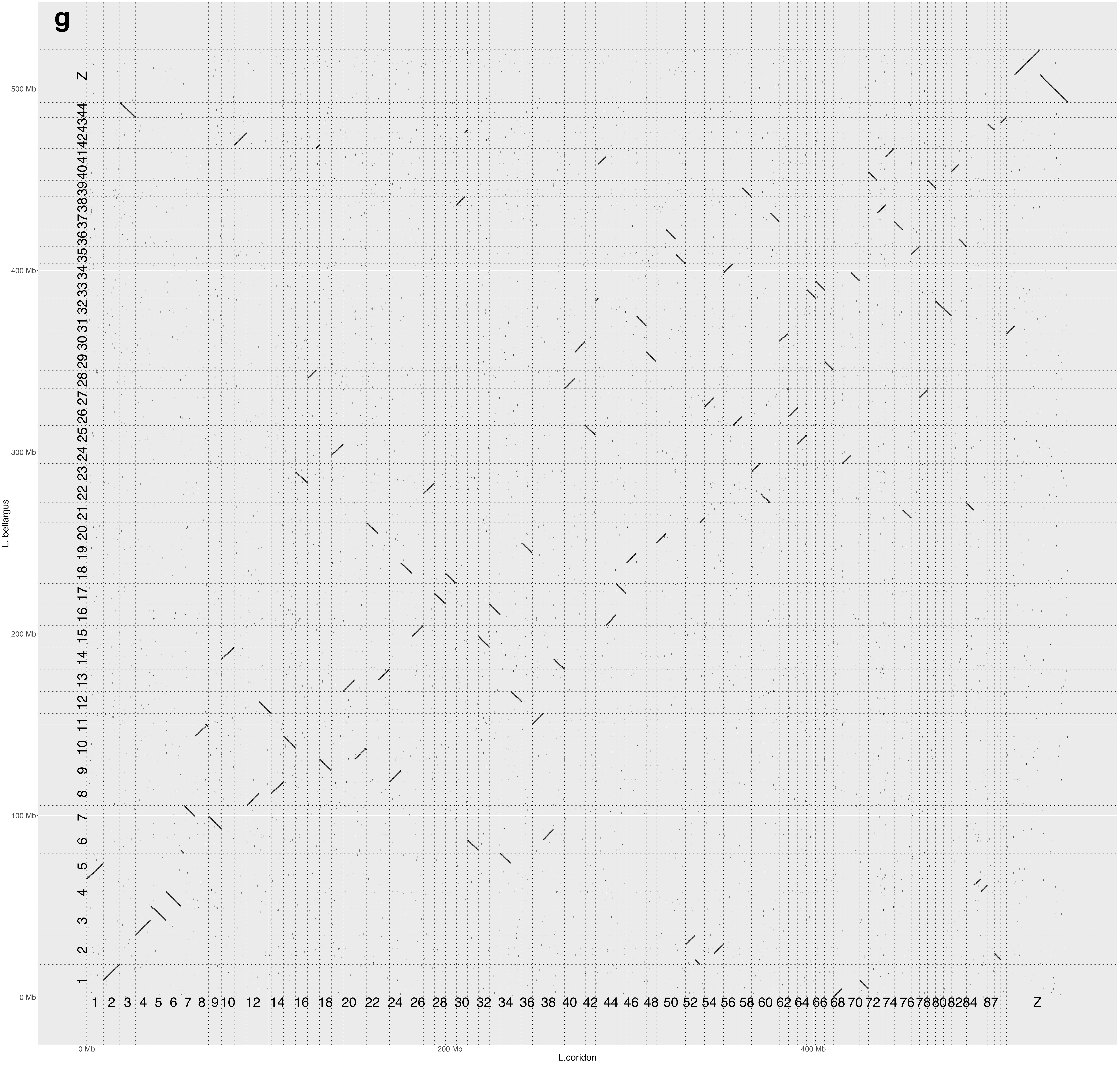
g
L. bellargus
L.coridon
0 Mb
100 Mb
200 Mb
300 Mb
400 Mb
500 Mb
Z
1 2 3 4 5 6 7 8 9 10 12 14 16 18 20 22 24 26 28 30 32 34 36 38 40 42 44 46 48 50 52 54 56 58 60 62 64 66 68 70 72 74 76 78 80 82 84 87 Z
